# MORONET: Multi-omics Integration via Graph Convolutional Networks for Biomedical Data Classification

**DOI:** 10.1101/2020.07.02.184705

**Authors:** Tongxin Wang, Wei Shao, Zhi Huang, Haixu Tang, Jie Zhang, Zhengming Ding, Kun Huang

**Author notes:** These authors contributed equally to this work (Tongxin Wang, Wei Shao).

## Abstract

To fully utilize the advances in omics technologies and achieve a more comprehensive understanding of human diseases, novel computational methods are required for integrative analysis for multiple types of omics data. We present a novel multi-omics integrative method named Multi-Omics gRaph cOnvolutional NETworks (MORONET) for biomedical classification. MORONET jointly explores omics-specific learning and cross-omics correlation learning for effective multi-omics data classification. We demonstrate that MORONET outperforms other state-of-the-art supervised multi-omics integrative analysis approaches from a wide range of biomedical classification applications using mRNA expression data, DNA methylation data, and miRNA expression data. Furthermore, MORONET is able to identify important biomarkers from different omics data types that are related with the investigated diseases.

## Introduction

The rapid advancement in high-throughput technologies have enabled the collection of various types of “omics” data at an unprecedented detailed level. Genome-wide data for different molecular processes, such as mRNA expression, DNA methylation, and microRNA (miRNA) expression, can be acquired for the same set of samples, resulting in multiple omics (multi-omics) data. While each omics technology itself can only capture part of the biological complexity in the investigated problem, the integration of multiple types of omics data is needed to provide a more comprehensive view of the underlying biological processes. For human diseases, existing studies have demonstrated that incorporating data from multiple omics technologies can improve the accuracy of predicting patient clinical outcomes performances comparing to using a single type of omics data^1–7^. Therefore, there is a strong motivation for integrative analysis methods to take advantage of the interactions and complementary information of multi-omics data.

A great number of methods have been proposed over the years for multi-omics data integration for a variety of problems. However, most existing research efforts focus on unsupervised multi-omics data integration without the additional information of sample labels. With the rapid development of personalized medicine, curated datasets with detailed annotations that characterize the phenotypes or traits of the samples are becoming more widely available. Therefore, there is an increasing interest in supervised multi-omics integration methods that can perform prediction on new samples, as well as identifying disease related biomarkers. Early attempts of supervised data integration for biomedical classification problems include direct concatenation of different types of omics data to learn the classification model^5^. Ensemble-based strategies have also been explored to integrate the predictions of the classifiers, each trained on one type of omics data individually^1^. However, these methods failed to consider the correlations among different omics data types and could be biased towards certain type of omics data. Recently, more supervised multi-omics integration methods have been proposed by exploiting the interactions across different omics data types. For example, van de Wiel *et al.*^6^ introduced an adaptive group-regularized ridge regression method that incorporated methylation microarray data and curated annotations of methylation probes for cervical cancer diagnostic classification. Singh *et al.*^4^ proposed DIABLO (Data Integration Analysis for Biomarker discovery using Latent cOmponents) by extending the sparse generalized canonical correlation analysis to a supervised framework, which could seek common information across multiple omics types while discriminating between different phenotypic groups.

With the continuous advancement of deep learning in various tasks, more and more multi-omics integration methods based on deep learning have been proposed to take advantage of the high learning capability and flexibility of deep neural networks^2, 8–10^. For example, Huang *et al.*^2^ integrated the features of mRNA expression and miRNA expression, along with additional clinical information at hidden layers for better prognosis prediction in breast cancer. However, these existing methods are based on fully-connected networks, which did not exploit the correlations between patients effectively through patient similarity networks. Moreover, while current deep learning based methods integrate different omics data at the input space^8, 10^ or the learned feature space^2, 9^, different omics data types could provide unique characteristics at the high-level label space. Therefore, it is crucial to utilize the correlations across different classes and different omics data types to further boost the learning performance.

To this end, we introduce MORONET, a multi-omics data analysis framework for classification tasks in biomedical applications. MORONET unifies omics-specific learning with multi-omics integrative classification at the label space. Specifically, MORONET utilizes Graph Convolutional Networks (GCN) for omics-specific learning. Comparing to the fully-connected neural networks, GCN can take advantage of both the omics features and the correlations among patients described by the patient similarity networks for better classification performance. While GCN has been utilized in unsupervised and semi-supervised settings, in this work, we extend the usage of GCN to supervised classification tasks on multi-omics data. MORONET also utilizes View Correlation Discovery Network (VCDN) for multi-omics integrative classification. VCDN can exploit the higher-level cross-omics correlations in the label space, as different omics data types could provide unique class-level distinctiveness. Therefore, it is crucial to explore the cross-omics label correlations to improve model performances on multi-omics data. While original form of VCDN was designed for samples with two views^11^, we further generalize it to accommodate three types of omics data: mRNA expression, DNA methylation, and miRNA expression. To the best of our knowledge, MORONET is the first supervised multi-omics integrative method for effective class prediction on new samples that not only utilizes GCN for omics data learning but also explores the cross-omics correlations at the label space. We demonstrate the capabilities and versatility of MORONET through a wide range of biomedical classification applications, including Alzheimer’s disease patient classification, tumor grade classification in low grade glioma, kidney cancer type classification, and breast invasive carcinoma subtype classification. We also showed the necessity of integrating multiple omics data types and the importance of combining both GCN and VCDN for multi-omics data classification through comprehensive ablation studies. Moreover, we demonstrated that MORONET is capable of identifying important omics signatures and biomarkers that are related to diseases of interests.

## Results

### Framework of MORONET

We introduce MORONET, a novel supervised multi-omics integration framework for a wide range of biomedical classification tasks (Figure 1). After pre-processing and feature pre-selection to remove noise and redundant features, we first use GCN to learn the classification task with each omics data type individually. Specifically, for each type of omics data, we construct a weighted patient similarity network using cosine similarity. Taking the input of both the omics features and the corresponding patient similarity network, GCN is trained for each omics data type to generate initial predictions of class labels. A major advantage of GCN is that they can take both the omics data and the correlations between patients for better prediction. Then, initial predictions generated by each omics-specific GCN were further utilized to produce the cross-omics discovery tensor, which reflects the cross-omics label correlations. Finally, the cross-omics discovery tensor is reshaped to a vector and forwarded to VCDN for final label prediction. VCDN can effectively integrate initial predictions from each omics-specific networks by explores the latent correlations across different omics data types in the higher-level label space. MORONET is an end-to-end model and omics-specific GCN and VCDN are trained alternatively until convergence. To this end, the final prediction is based on both effective omics-specific predictions generated by GCN and the learned cross-omics label-correlation knowledge generated by VCDN. To the best of our knowledge, MORONET is the first method to combine GCN and VCDN for effective multi-omics integration in biomedical data classification applications.

**Figure 1.**
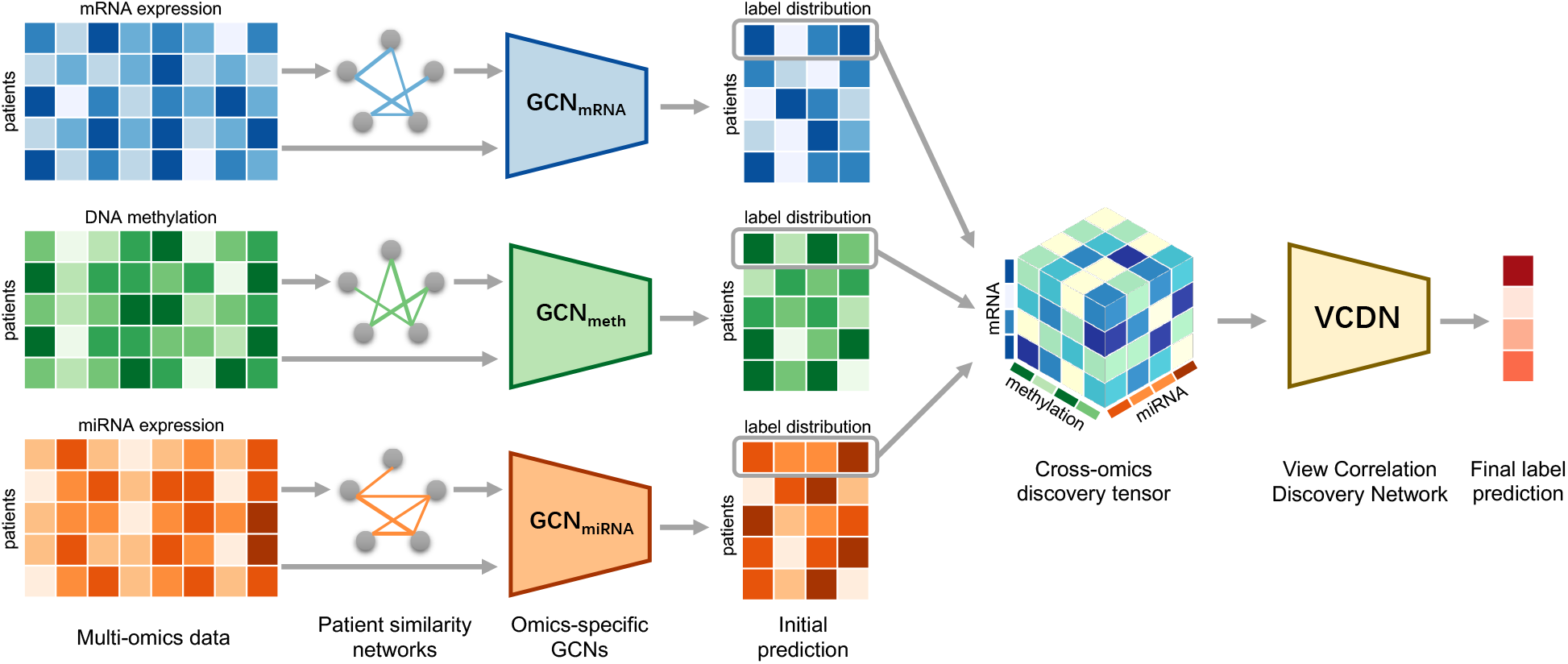
Illustration of MORONET. MORONET combines GCN for multi-omics specific learning and VCDN for multi-omics integration. For concise illustration, an example of one patient is chosen to demonstrate the VCDN component for multi-omics integration after omics-specific learning. Pre-processing is first performed on each omics data type to remove noise and redundant features. Omics-specific GCN learns class prediction using omics features and the corresponding patient similarity network generated from the omics data. Cross-omics discovery tensor is calculated from initial predictions from GCN and forwarded to VCDN for final prediction. MORONET is an end-to-end model and all networks are trained jointly.

### Datasets

To demonstrate the effectiveness of MORONET, we applied the proposed method on four different biomedical classification tasks using four different datasets: ROSMAP for Alzheimer’s disease (AD) patients versus normal control (NC) classification, LGG for grade classification in Low Grade Glioma (LGG), KIPAN for kidney cancer type classification, and BRCA for Breast Invasive Carcinoma (BRCA) subtype classification. Among these tasks, kidney cancer type classification is the simplest task and serves more as a proof-of-concept experiment for multi-class applications, since the differences among chromophobe renal cell carcinoma (KICH), clear renal cell carcinoma (KIRC), and papillary renal cell carcinoma (KIRP) can be clearly observed in the omics data. The details of the datasets are listed in Table 1. Specifically, ROSMAP is composed of ROS and MAP, both are longitudinal clinical-pathologic cohort studies of AD from Rush University^12, 13^. The ROSMAP dataset was acquired from AMP-AD Knowledge Portal (https://adknowledgeportal.synapse.org/)^14^. AD patients and normal control subjects were selected for the classification task in our experiment. Omics data of LGG, KIPAN, and BRCA, as well as the grade information of LGG patients were acquired from The Cancer Genome Atlas Program (TCGA) through Broad GDAC Firehose (https://gdac.broadinstitute.org/). PAM50 breast cancer subtypes of TCGA BRCA patients^15, 16^ were acquired through TCGAbiolinks^17^. Three types of omics data (*i.e.* mRNA expression data, DNA methylation data, and miRNA expression data) were used for classification to provide comprehensive and complementary information about the diseases. Only subjects with matched mRNA expression, DNA methylation, and miRNA expression data were included in our study.

**Table 1.** Summary of datasets

| Dataset | Categories | Number of features |  |  |
| --- | --- | --- | --- | --- |
|  |  | mRNA | DNA methylation | miRNA |
| ROSMAP | NC: 169, AD: 182 | 55889 | 23788 | 309 |
| LGG | Grade 2: 246, Grade 3: 264 | 20531 | 20114 | 548 |
| KIPAN | KICH: 66, KIRC: 318, KIRP: 274 | 20531 | 20111 | 445 |
| BRCA | Normal: 115, Basal: 131, Her2: 46, LumA: 436, LumB: 147 | 20531 | 20106 | 503 |

### Multi-omics classification performance evaluation

In this section, we compared the classification performance of MORONET with existing supervised multi-omics integration algorithms, as well as performing comprehensive ablation studies to demonstrate the necessity of different components in MORONET. To compare the effectiveness of different multi-omics integration methods, we randomly selected 30% of the samples in a dataset as the test set and the remaining 70% of the samples as the training set. The test set was constructed by preserving the class distribution in the original dataset. To evaluate the performance of the compared methods, we calculated accuracy (ACC) and F1 score (F1) of the classification results. Weighted average of F1 score by support for each label was used to account for label imbalance. For binary classification tasks, Area Under the Receiver Operating Characteristic Curve (AUC) was also reported. We evaluated all the methods on five different randomly generated training and test splits and the mean and standard deviation of the evaluation metrics across these five experiments were reported.

### MORONET outperformed existing supervised multi-omics integration methods in various classification tasks

We compared the classification performance of MORONET with the following eight existing classification algorithms: 1) K-nearest neighbor classifier (KNN): label predictions were made by voting of k-nearest neighbors in the training data. We set *k* = 5 in all experiments. 2) Support vector machine classifier (SVM). 3) Linear regression trained with L1 regularization (LASSO). In LASSO, an individual model was trained to predict the probability of each class, and the class predicted with the highest probability was selected as the final class label prediction for the entire model. 4) Random forest classifier (RF). 5) Fully-connected neural network classifier (NN): deep fully-connected neural network with three layers trained with cross entropy loss. 6) Adaptive group-regularized ridge regression (GRridge)^6^: Implementation in the GRridge R package was used. 7) block_plsda: multi-omics integration with projection to latent structures models with discriminant analysis. Block_plsda integrates multiple types of omics data measured on the same samples to classify a discrete outcome. Block_plsda is one of the supervised analysis methods included in DIABLO^4^. 8) block_splsda: block_plsda with additional sparse regularization, which can select relevant features from each dataset. It is also a supervised analysis method within DIABLO. Implementations in the mixOmics R package^18^ were used for block_plsda and block_splsda. Block_plsda and block_splsda represent the state-of-the-art approaches for supervised multi-omics integration and classification. KNN, SVM, LASSO, and NN were trained with direct concatenation of multi-omics data as input. All methods were trained with the same pre-processed data. The classification results for ROSMAP, LGG, KIPAN, and BRCA are shown in Tables 2–5, respectively.

**Table 2.** Classification results on ROSMAP dataset

| Method | ACC | F1 | AUC |
| --- | --- | --- | --- |
| KNN | 66.79 ± 5.70 | 66.76 ± 5.67 | 71.38 ± 6.71 |
| SVM | 77.92 ± 0.75 | 77.92 ± 0.76 | 77.97 ± 0.78 |
| LASSO | 70.19 ± 3.51 | 70.02 ± 3.41 | 78.13 ± 3.86 |
| RF | 70.38 ± 3.41 | 70.17 ± 3.19 | 76.66 ± 2.66 |
| NN | 68.30 ± 5.71 | 68.21 ± 5.70 | 74.20 ± 4.88 |
| GRridge | 76.79 ± 2.64 | 76.76 ± 2.65 | 84.63 ± 3.97 |
| block_plsda | 75.28 ± 1.39 | 75.23 ± 1.36 | 83.92 ± 1.54 |
| block_splsda | 76.42 ± 2.46 | 76.38 ± 2.46 | 83.89 ± 2.58 |
| NN_NN | 79.62 ± 2.64 | 79.53 ± 2.69 | 84.61 ± 2.13 |
| NN_VCDN | 79.25 ± 2.98 | 79.21 ± 3.00 | 83.91 ± 3.67 |
| GCN_NN | 81.32 ± 3.29 | 81.31 ± 3.28 | 86.80 ± 2.17 |
| MORONET | **82.45 ± 2.20** | **82.41 ± 2.21** | **87.34 ± 2.18** |

**Table 3.** Classification results on LGG dataset

| Method | ACC | F1 | AUC |
| --- | --- | --- | --- |
| KNN | 73.07 ± 3.24 | 73.07 ± 3.24 | 80.06 ± 3.10 |
| SVM | 74.38 ± 2.39 | 74.37 ± 2.38 | 74.36 ± 2.36 |
| LASSO | 76.47 ± 1.55 | 76.46 ± 1.56 | 83.66 ± 2.52 |
| RF | 74.77 ± 2.50 | 74.71 ± 2.48 | 81.77 ± 1.52 |
| NN | 72.55 ± 3.28 | 72.53 ± 3.27 | 78.76 ± 2.80 |
| GRridge | 70.59 ± 1.31 | 70.53 ± 1.25 | 77.42 ± 1.72 |
| block_plsda | 73.73 ± 3.39 | 73.56 ± 3.45 | 81.65 ± 2.33 |
| block_splsda | 78.95 ± 2.24 | 78.90 ± 2.24 | **85.86 ± 1.82** |
| NN_NN | 73.73 ± 2.16 | 73.58 ± 2.16 | 78.78 ± 2.52 |
| NN_VCDN | 72.94 ± 3.45 | 72.74 ± 3.45 | 77.61 ± 4.10 |
| GCN_NN | 78.95 ± 1.72 | 78.91 ± 1.71 | 85.84 ± 1.80 |
| MORONET | **79.48 ± 2.36** | **79.46 ± 2.37** | 85.43 ± 2.29 |

**Table 4.** Classification results on KIPAN dataset

| Method | ACC | F1 |
| --- | --- | --- |
| KNN | 96.77 ± 0.94 | 96.76 ± 0.94 |
| SVM | 99.09 ± 0.87 | 99.09 ± 0.87 |
| LASSO | 97.17 ± 0.40 | 97.16 ± 0.41 |
| RF | 97.88 ± 0.59 | 97.87 ± 0.59 |
| NN | 98.59 ± 0.20 | 98.58 ± 0.21 |
| GRridge | 99.49 ± 0.32 | 99.49 ± 0.32 |
| block_plsda | 95.25 ± 0.82 | 95.23 ± 0.82 |
| block_splsda | 94.95 ± 0.45 | 94.93 ± 0.45 |
| NN_NN | 99.80 ± 0.40 | 99.80 ± 0.40 |
| NN_VCDN | 99.60 ± 0.59 | 99.59 ± 0.59 |
| GCN_NN | 99.70 ± 0.25 | 99.69 ± 0.25 |
| MORONET | **99.80 ± 0.25** | **99.80 ± 0.25** |

**Table 5.**
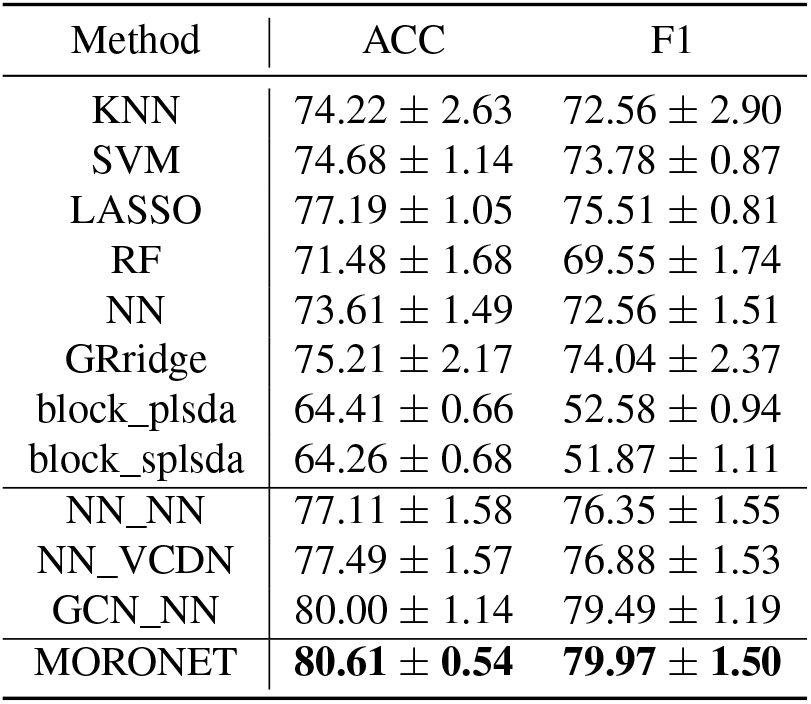
Classification results on BRCA dataset

| Method | ACC | F1 |
| --- | --- | --- |
| KNN | 74.22 ± 2.63 | 72.56 ± 2.90 |
| SVM | 74.68 ± 1.14 | 73.78 ± 0.87 |
| LASSO | 77.19 ± 1.05 | 75.51 ± 0.81 |
| RF | 71.48 ± 1.68 | 69.55 ± 1.74 |
| NN | 73.61 ± 1.49 | 72.56 ± 1.51 |
| GRridge | 75.21 ± 2.17 | 74.04 ± 2.37 |
| block_plsda | 64.41 ± 0.66 | 52.58 ± 0.94 |
| block_splsda | 64.26 ± 0.68 | 51.87 ± 1.11 |
| NN_NN | 77.11 ± 1.58 | 76.35 ± 1.55 |
| NN_VCDN | 77.49 ± 1.57 | 76.88 ± 1.53 |
| GCN_NN | 80.00 ± 1.14 | 79.49 ± 1.19 |
| MORONET | **80.61 ± 0.54** | **79.97 ± 1.50** |

From Tables 2–5, we observed that MORONET outperformed the compared multi-omics integration methods in most classification tasks. The only exception was in LGG grade classification, where block_splsda yielded slightly higher AUC score than MORONET. However, MORONET achieved better performance in LGG grade classification when evaluated through ACC and F1 score. Moreover, MORONET consistently outperformed the state-of-the-art supervised multi-omics integration methods (block_plsda and block_splsda) in all the other tasks, which demonstrated the superiority of multi-omics data classification capability by combining GCN for omics-specific learning and VCDN for multi-omics integration. Comparing with existing methods, while MORONET yielded the best results in simple tasks like kidney cancer type classification, its significant advantages were demonstrated in more difficult applications such as AD patient classification and BRCA subtype classification, demonstrating the superior learning capability of MORONET. Interestingly, although deep learning based methods have shown great promises in classification applications, the deep learning based method NN did not show obvious improvements comparing with other shallow methods. This suggests that proper design of deep learning algorithms specific to supervised multi-omics integration applications is required to achieve superior classification performance.

### MORONET outperformed its variations in various classification tasks

MORONET combines omics-specific learning via GCN with cross-omics correlation learning using VCDN for effective multi-omics data classification. GCN can learn from both features and the graph structure of the training data, which explicitly takes advantage of the correlations among the training samples comparing to the commonly used fully-connected neural networks. VCDN is designed to learn the higher-level intraview and cross-view correlations in the label space in multiview learning problems, which can boost the performance of multi-omics integration. To demonstrate the necessity of GCN and VCDN for effective multi-omics data classification, we performed extensive ablation studies of our proposed method where three additional variations of MORONET were compared. 1) NN_NN: Fully-connected neural networks with the same number of layers and the same dimensions of hidden layers as MORONET was used for omics-specific classification learning. A neural network with the same structure as VCDN was used for multi-omics integration. However, instead of constructing the cross-omics discovery tensor, label distribution from each omics data type was directly concatenated to a longer vector as the input of the multi-omics integration network. 2) NN_VCDN: The omics-specific classification component was the same as NN_NN without utilizing GCN. The multi-omics integration component utilized VCDN, which was the same as MORONET. 3) GCN_NN: The omics-specific classification component utilized GCN, which was the same as MORONET. The multi-omics integration part was the same as NN_NN without VCDN.

From Tables 2–5, we observed that MORONET outperformed its three variations in ablation studies for most classification tasks. In the LGG grade classification, GCN_NN yielded slightly higher AUC score than MORONET, while MORONET achieved better ACC and F1 score. One possible reason of the similar performance between GCN_NN and MORONET in this task could be due to the fact that it was a binary classification problem, where the contribution of cross-view correlation in the label space might be limited as the number of distinct labels were limited to two. Specifically, GCN_NN and MORONET shared the same structure for multi-omics integration component except that the input dimensions were different. For binary classification problems, the input dimension for the multi-omics integration component in GCN_NN was 2 × 3 = 6, while the input dimension for the same component in MORONET was 2^3^ = 8. While VCDN can effectively utilize the cross-view correlation in the label space, such advantage can be limited when the number of distinct labels was small. Nevertheless, exploring cross-view correlations was still essential to multi-omics classification as we observed that MORONET outperformed GCN_NN in most binary classification tasks and in all multi-class classification tasks under different evaluation metrics. Moreover, we observed that MORONET consistently outperformed NN_VCDN in all classification tasks, which demonstrated that the classification performance can be improved by not only learning from the omics features, but also learning from the patient correlations and the topological structures of the training samples through GCN. Another interesting observation was that while MORONET consistently outperformed NN_NN, both NN_VCDN and GCN_NN failed to consistently outperformed NN_NN in all the tasks, which suggested that GCN and VCDN need to be combined and trained jointly in order to achieve superior results for multi-omics classification tasks.

To further demonstrate the necessity of integrating multiple types of omics data to boost the classification performance in biomedical applications, we compared the final classification performance of MORONET with the classification results of each omics data type produced by omics-specific GCN before integration. The results are shown in Figure 2. From Figure 2, we observed that by exploring the cross-omics label correlations through VCDN, the classification performance was consistently improved by integrating classification results of multiple omics data types. Specifically, for classification on ROSMAP, LGG, and BRCA, results produced by MORONET are significantly better than any of the single-omics classification results. For KIPAN, while results of MORONET were slightly better than only using DNA methylation data, MORONET was able to produce more consistent results with the standard deviation of ACC and F1 score greatly reduced through multi-omics integration.

**Figure 2.**
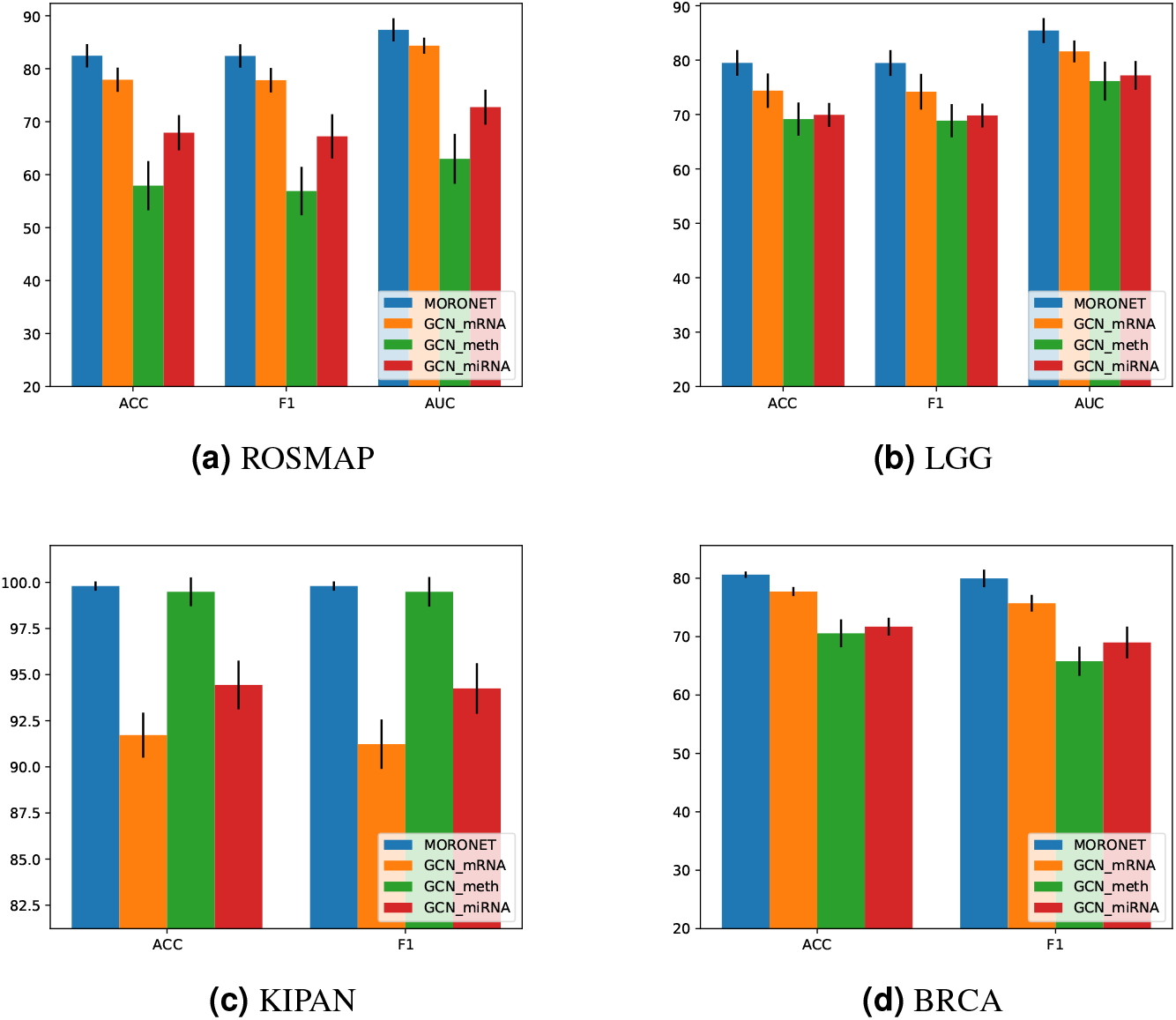
Performance comparison of multi-omics data classification via MORONET and single-omics data classification via GCN. GCN_mRNA, GCN_meth, and GCN_miRNA refers to single-omics data classification via GCN with mRNA expression data, DNA methylation data, and miRNA expression data, respectively.

### Important biomarkers identified by MORONET

Identifying biomarkers is essential to interpreting the results in biomedical applications and understanding the underlying biology of the problems. There have been extensive studies of interpreting feature importance for neural networks over the years. Since the input of MORONET is scaled to [0, 1] during pre-processing, we can remove the signal from a feature by setting it to zero. Therefore, the importance of a feature to the classification task can be measured by the extent of performance drop after the feature is set to zero. This approach has been widely adopted for feature importance ranking and feature selection in neural networks^2, 19–21^. Based on this approach, we analyzed the contribution of each feature in all types of omics data by assigning the feature as zero and calculated the classification performance decrease on the test set comparing to using all the features. Features with the largest performance drop were considered to be the most important ones. We used AUC to measure the performance drop for binary classification tasks and ACC for multi-class classification tasks. We selected the top 50 important features for each omics data type, and the features that were selected in more than three out of five repeated experiments in a dataset were reported. As mentioned in the previous section, the KIPAN dataset served as a proof-of-concept experiment for multi-class applications, and therefore was excluded from detailed biomarker identification analysis. For ROSMAP, LGG, and BRCA dataset, the identified mRNA expression, DNA methylation, and miRNA expression features are shown in in Tables 6–8 for further discussion.

**Table 6.**
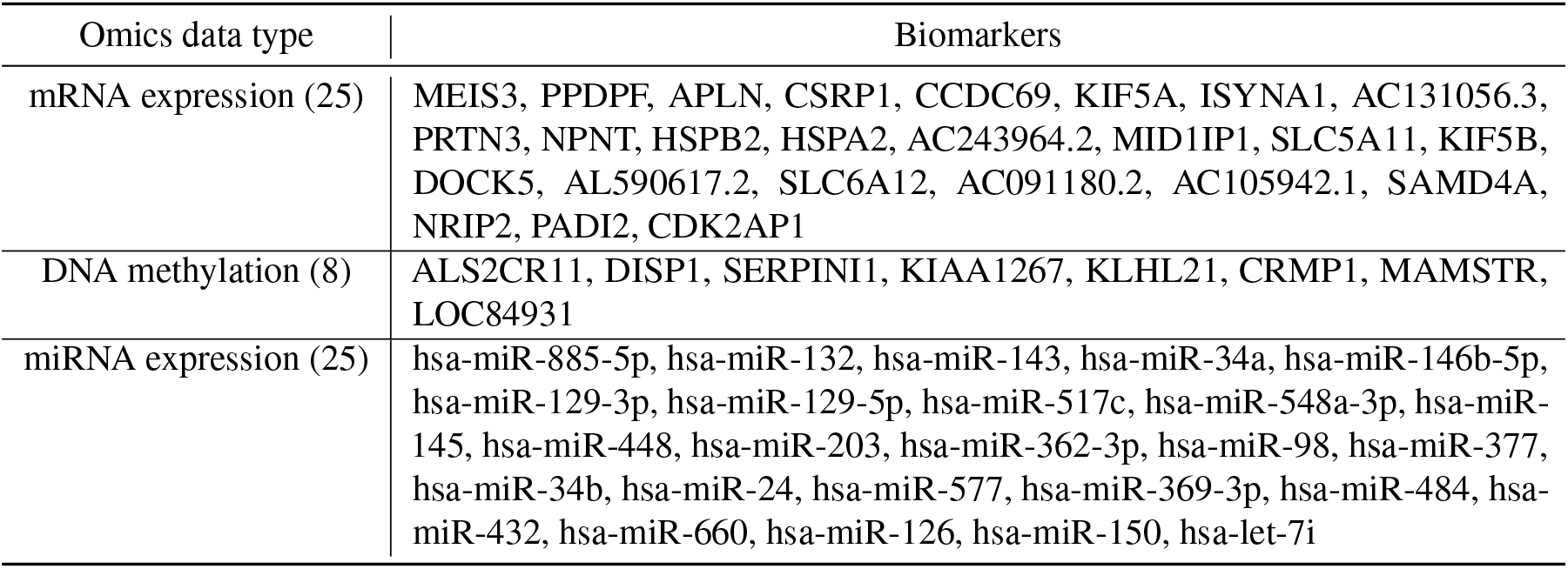
Important omics biomarkers identified in ROSMAP dataset

### MORONET identified biomarkers related to Alzheimer’s disease from ROSMAP dataset

For AD patient classification, 25 mRNA expression features, 8 DNA methylation features, and 25 miRNA expression features were identified by MORONET (Table 6). Specifically, genes identified by the mRNA expression data and genes corresponding to the identified DNA methylation features have been found associated with AD. For example, for identified mRNA expression features, Wang *et al.*^22^ found that the loss of kinesin-1 KIF5A isoform was a primary neuronal pathology in AD. They discovered that KIF5A deficiency was a novel mechanism of AD-relevant axonal mitochondrial traffic abnormalities and suggested a potential therapeutic treatment of AD by protecting the KIF5A function. Brock *et al.*^23^ found that PRTN3 expression level was significantly decreased in the occipital lobe of the brains with AD. Higher expression level of HSPB2 was also discovered to be associated with faster cognitive decline in Alzheimer’s dementia^24^. Petyuk *et al.*^25^ identified HSPA2 as an important regulator of late-onset Alzheimer’s disease processes. Moreover, microRNAs identified by the proposed algorithm were also found to be related to AD. For example, miR-885-5p and miR-143 were significantly down-regulated in the sera from the AD patients compared to negative controls^26, 27^. The abnormal expression of miR-34a was suggested to contribute to the progression of AD by affecting the expression level of BCL2^28^. Lau *et al.*^29^ identified that decrease of miR-132 expression in AD brains was most notable in neurons displaying Tau hyper-phosphorylation and suggested that miR-132 contributes to AD progression. Expression level of miR-132 was also reported to be correlated with insoluble tau and cognitive impairment^30^.

### MORONET identified biomarkers related to tumor grade in LGG from LGG dataset

For LGG grade classification, 24 mRNA expression features, 16 DNA methylation features, and 24 miRNA expression features were identified by MORONET (Table 7). For genes of the identified mRNA expression features and genes corresponding to the identified DNA methylation features, we applied ToppGene Suite^31^ for gene set functional enrichment analysis to determine if genes identified by MORONET are biologically meaningful. ToppGene finds biological annotations such as Gene Ontology (GO) items that are significant in a set of genes and multiple-testing corrections are applied to the reported p values. For genes identified in mRNA expression features, they are significantly enriched with GO biological process terms such as nuclear division (GO:0000280, *p* = 6.149*E* 8), DNA repair (GO:0006281, *p* = 6.149*E* 8), cell cycle (GO:0007049, *p* = 6.149*E* 8), and DNA metabolic process (GO:0006259, *p* = 9.262*E* 8). For genes identified in DNA methylation features, significantly enriched biological processes included keratinocyte differentiation (GO:0030216, *p* = 5.608*E* 3) and epidermal cell differentiation (GO:0009913, *p* = 8.185*E* 3). These biological processes are highly related to the development and the aggressiveness of cancer. Cancer is a disease of inappropriate cell proliferation, which results from the failure of the proper regulation of the cell cycle machinery^32^. Moreover, cell differentiation is strongly associated with the aggressiveness of the cancer, as poorly differentiated or undifferentiated cancer cells look and behave very differently from normal cells. Tumours with poorly differentiated or undifferentiated cancer cells tend to grow more aggressively with a higher risk of metastasis than tumours with well-differentiated cancer cells. Cell differentiation and the speed of cell proliferation and division are the most important factors in tumor grading, which is also consistent with the classification task of LGG tumor grades.

**Table 7.** Important omics biomarkers identified in LGG dataset

| Omics data type | Biomarkers |
| --- | --- |
| mRNA expression (24) | ITPRIPL1, AURKB, MCM2, CDC45, TEF, FANCA, DTL, BRIP1, MKI67, POC1A, PLAT, GSG2, ZNF367, SSTR1, POLQ, TACC3, FANCC, TTK, TK1, KIAA0101, RAD51, CHAF1A, IGFBP2, KIF4A |
| DNA methylation (16) | IL32, IQCF5, OR51A7, TGM3, BASE, SPRR1B, SPRR2D, OR8G1, FAM71F1, MIR298, C3orf22, INHBE, AQP10, SFTPB, OR13C3, REG3G |
| miRNA expression (24) | hsa-mir-383, hsa-mir-10b, hsa-mir-329-1, hsa-mir-491, hsa-mir-508, hsa-mir-29c, hsa-mir-184, hsa-mir-21, hsa-mir-128-2, hsa-mir-128-1, hsa-mir-218-2, hsa-mir-1296, hsa-mir-129-1, hsa-mir-27a, hsa-mir-488, hsa-mir-9-3, hsa-mir-193a, hsa-mir-767, hsa-mir-770, hsa-mir-103-1, hsa-mir-876, hsa-mir-885, hsa-mir-196b, hsa-mir-23a |

Besides biomarkers related to tumor grading, genes related to glioma were also identified by MORONET. For example, MKI67 and IGFBP2 were demonstrated as important biomarkers for determining the prognosis of glioma patients^33, 34^. MKI67 is one of the most widely used malignancy markers in cancer pathology^35, 36^ while IGFBP2 also have critical contribution to glioma development. Expression of IGFBP2 in gliomas was found correlated to the histological grade of the tumor^37^. Wang *et al.*^38^ also discovered that IGFBP2 could contributes to glioma progression and tumor cell invasion in part by enhancing MMP-2 gene transcription. Identified miRNA expression features have also been demonstrated to be involved in the regulation of glioma progression. For example, miR-383 was suggested to play the role of tumor suppressor in glioma cells by downregulating CCND1 expression^39^. Downregulation of miR-383 was suggested to enhance glioma cell invasive ability by participating in the regulation of constitutive IGF1R signaling activation^40^. Moreover, miR-10b was also found associated with glioma pathological grade and malignancy, as well as promoting glioma cell invasion by targeting HOXD10^41^.

### MORONET identified biomarkers related to BRCA subtypes from BRCA dataset

For BRCA PAM50 subtype classification, 35 mRNA expression features, 25 DNA methylation features, and 31 miRNA expression features were identified (Table 8). A lot of well-known breast cancer genes were identified by mRNA expression data. For example, enrichment analysis results produced by ToppGene showed that identified genes by mRNA expression data were significantly associated with breast adenocarcinoma (*p* = 4.665*E* 3), including 5 well-known breast cancer genes (AR^42, 43^, ERBB4^44^, FOXA1^42, 45–47^, BCL2^48, 49^, and TFF1^50^) according to the DisGeNET knowledge platform^51^. Specifically, Ni *et al.*^42^ found AR highly expressed in ER–/HER2+ breast tumors and also identified important collaboration between AR and FOXA1 in transcriptional activation of AR target genes in ER–/HER2+ breast cancer cells. High expression levels of FOXA1 were observed in ER+ breast cancers^45^. FOXA1 was also found correlated with LumA breast cancer and was identified as a significant prognosis predictor in patients with ER+ tumors^46, 47^. Genes identified by mRNA expression data were also significantly enriched in molecular functions such as sequence-specific DNA binding (GO:0043565, *p* = 4.422*E* 3), transcription regulatory region DNA binding (GO:0044212, *p* = 4.422*E* 3), and regulatory region nucleic acid binding (GO:0001067, *p* = 4.422*E* 3), which is consistent with the important role of transcriptional regulation in different breast cancer subtypes. Moreover, genes related to BRCA were also identified by DNA methylation data. For example, higher expression levels of LRRC25 were reported to be suggestively associated with increased risk of breast cancer^52^. SOSTDC1 expression was found reduced in breast cancer compared to normal breast tissue, and high SOSTDC1 expression levels were found correlated with increased survival in breast cancer patients^53^. BRCA-related miRNAs were also identified by MORONET. For example, Yang *et al.* showed that miR-223 can promote the invasion of breast cancer cells via the Mef2c-*β* - catenin pathway^54^. Expression levels of miR-204 were found to have lower expression levels in breast cancer tissues than in the adjacent normal breast tissues^55^. Shen *et al.*^56^ discovered that miR-204 can regulate the biological behavior of breast cancer cells through directly targeting FOXA1. Moreover, for breast cancer PAM50 subtype related miRNA differential expression analysis, miR-223 was found downregulated in LumB breast cancers and miR-204 was found upregulated in normal breast cancers^57^.

**Table 8.**
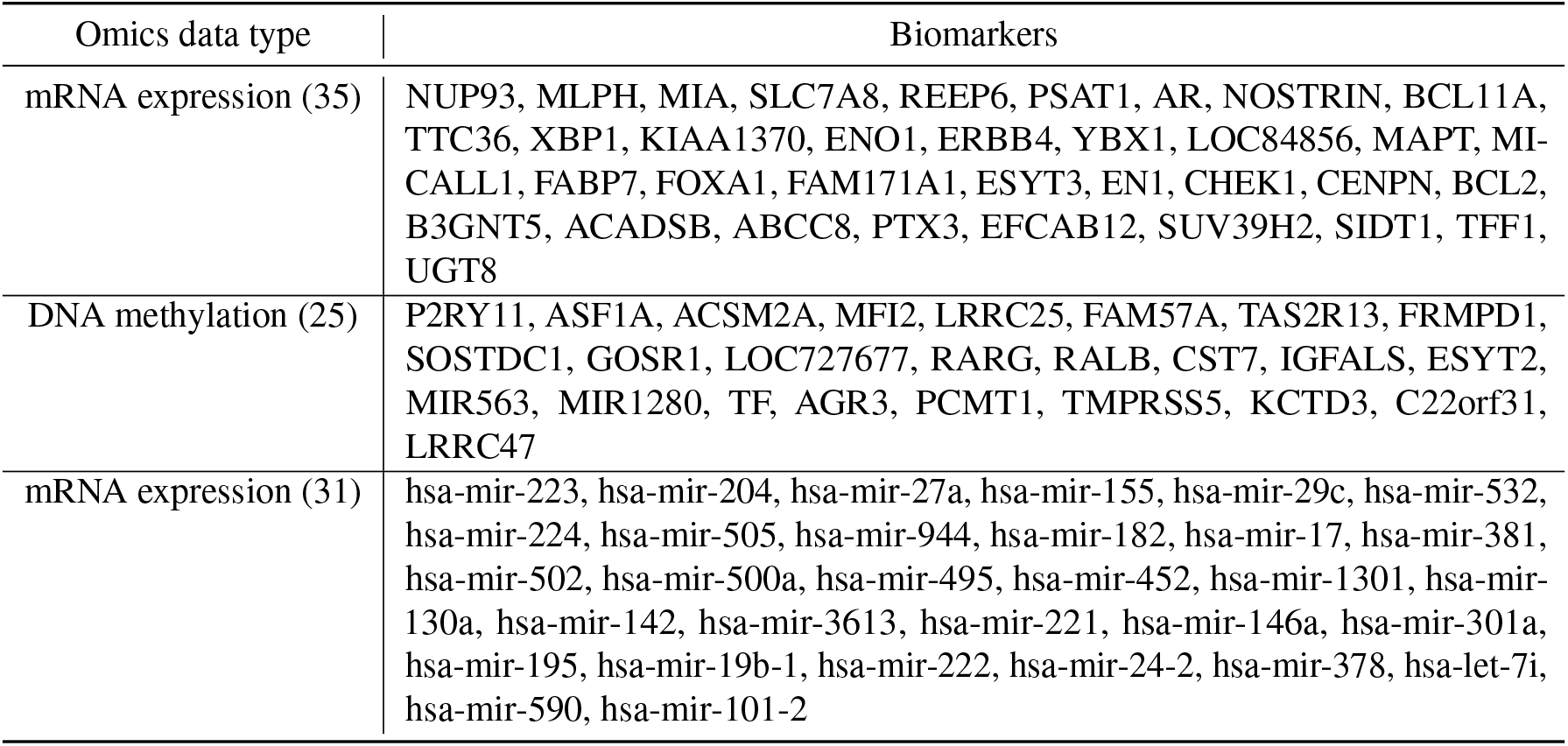
Important omics biomarkers identified in BRCA dataset

## Discussion

The rapid advancement of omics technologies has enabled personalized medicine at molecular level with unprecedented details. With the ability of measuring the same set of samples with multiple omics technologies, integration of multiple omics data types is needed to provide a more comprehensive view of human diseases as each technology itself can only characterize part of the underlying biology. Previously, labeled biomedical data have been scarce as manually collecting and annotating data are highly expensive and time consuming. Consequently, most existing multi-omics integration methods focus on unsupervised methods without additional phenotypic information and extracting biological insights from the identified clusters of samples. However, thanks to the rapid development of omics technologies and personalized medicine, labeled omics datasets with detailed annotations are becoming available at an unprecedented volume and speed. Therefore, it has become more and more important to take advantage of these labeled omics data to better predict essential phenotypes or traits (*e.g.*, disease diagnosis, grading of tumors, and cancer subtypes) on new samples.

To this end, we propose MORONET, a novel supervised multi-omics integration method for biomedical classification tasks based on deep multi-view learning. We consider each omics data type as a view of the samples. We utilized GCN for omics-specific learning and VCDN for multi-omics integration at the high-level label space. We demonstrated that MORONET could outperform state-of-the-art supervised multi-omics integration methods in a variety of biomedical classification applications, such as AD patient classification, tumor grade classification in LGG, kidney cancer type classification, and BRCA PAM50 subtype classification.

Through rigorous ablation studies, we demonstrated that both GCN and VCDN are essential to effective multi-omics data classification. Comparing to fully-connected networks, GCN can utilize both the features and the graphical structures of the data. Such graphical structure is important for biological data given the extensive interactions among genes and molecules. In MORONET, by constructing the patient similarity networks from omics data, both the omics features and the correlation between samples can be explicitly and simultaneously utilized through GCN. While commonly-used fully-connected networks can only be trained on structured data, GCN can also generalize neural networks to work on arbitrarily structured graphs. This suggests that our GCN-based method is flexible and can be generalized to include more types of information to boost the classification performance in the future. Comparing with traditional applications of GCN, which are either learning embeddings of graphs in an unsupervised fashion or learning to propagate labels from labeled samples to unlabeled samples in the graph in an semi-supervised fashion, MORONET further extend the use of GCN to supervised learning for better classification of multi-omics data on new samples. To the best of our knowledge, our method is one of the first methods to explore GCN in supervised multi-omics integration for classification tasks. We also demonstrated that VCDN can effectively classify multi-omics data by integrating the omics-specific classification produced by GCN at the label space. Comparing to directly concatenating class distributions predicted by GCN, VCDN can effectively explore the cross-omics correlations in the label space to boost the multi-omics data classification performance. Comparing to the original application of VCDN on human action recognition tasks^11^, which only considered data with two views, we further extended it to accommodate multiple omics data types.

One important hyper-parameter in MORONET is *k*, which determines the threshold of affinity values adaptively when constructing the weighted patient similarity networks for omics-specific GCNs. In our applications, *k* represents the average number of edges per patient that are retained in the patient similarity networks except self loops. Patient similarity networks that faithfully capture the interactions between patients can boost the performance of GCN by providing additional information of patient correlations. However, if *k* is set too small, the patient similarity network becomes too sparse and some important patient interactions could be missed. On the other hand, if *k* is too large, the patient similarity network becomes too dense and noise or artifacts of patient correlations might be included. Therefore, choosing a proper *k* value is important to the performance of MORONET. However, a proper choice of *k* depends on the topological structure of data, which varies from dataset to dataset. In our experiments, *k* is determined through cross-validation on the training data. To further demonstrate the effects of hyper-parameter *k* on the performance of MORONET in both binary and multi-class classification tasks, we trained MORONET under a wide range of *k* values using the ROSMAP dataset and BRCA dataset. Figure 3 shows the performance of MORONET when *k* varies from 1 to 10, where the dashed lines represent the results from the best performed existing multi-omics integration methods. From 3, we observed that the hyper-parameter *k* did influence the classification performance of MORONET as the performance fluctuates with the changes of *k*. However, MORONET was robust to different values of *k* as it consistently outperformed existing methods under a wide range of *k* values.

**Figure 3.**
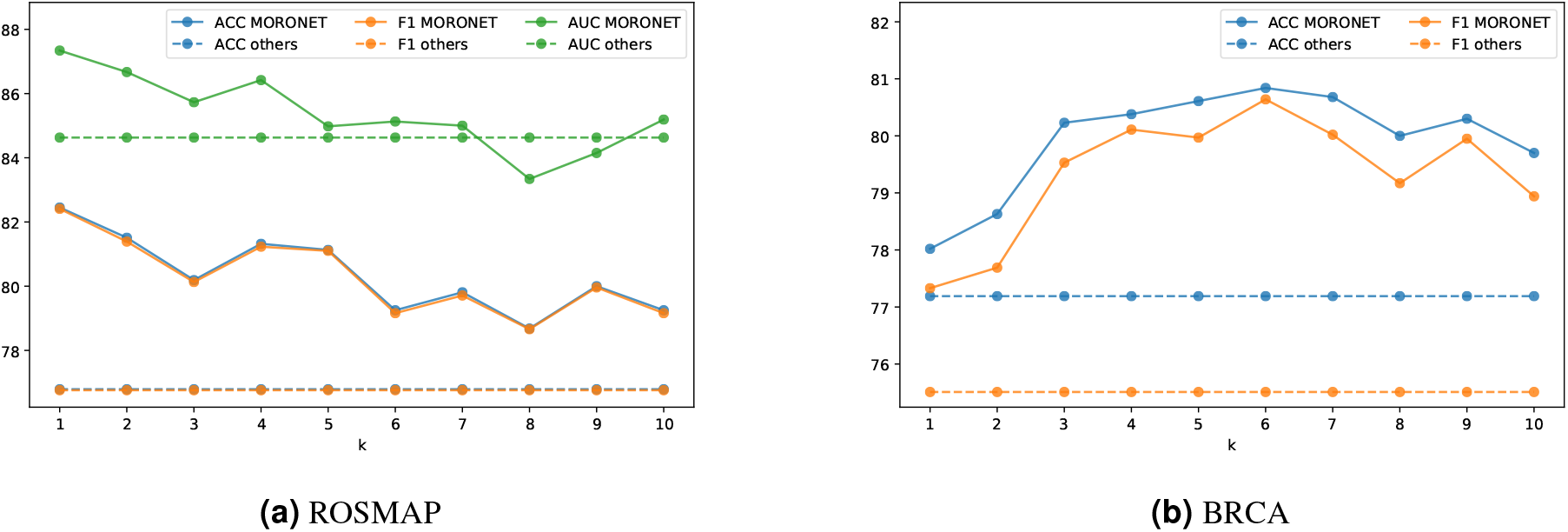
Performance of MORONET under different values of hyper-parameter *k*. The dashed lines represent the results from the best performed existing multi-omics integration methods. MORONET consistently outperformed existing methods under a wide range of *k* values.

## Conclusion

Integration of multiple types of omics data is essential to achieve a more comprehensive understanding of the underlying biology in diseases. With the improvement in omics technologies and the wide availability of well-annotated omics datasets, there is a need for classification methods utilizing multi-omics data to better predict important phenotypes or traits on new samples. In this paper we introduce MORONET, a novel classification method for multi-omics data based on deep multi-view learning. MORONET uses GCN for omics-specific classification learning, which can utilize both the omics features and the correlations between samples. MORONET uses VCDN to explores cross-omics correlations in the label space for effective multi-omics integration. MORONET demonstrated significant improvements in a wide range of multi-omics classification tasks for human diseases, such as AD patient classification, tumor grade classification in LGG, kidney cancer type classification, and BRCA PAM50 subtype classification, comparing to single-omics classification, existing state-of-the-art multi-omics classification methods, and its own variations in ablation studies. MORONET also effectively identified meaningful genomic features in each omics data type that showed strong association with the diseases of interest. Therefore MORONET is an innovative deep learning based multi-omics classification algorithm with both superior performance and good interpretability.

## Methods

### Method overview

MORONET is a novel framework for a variety of biomedical classification tasks utilizing multi-omics data. The workflow of MORONET can be summarized into three components (Figure 1): (1) Pre-processing. Pre-processing and feature pre-selection were performed on each omics data types individually to remove noise, artifacts, and redundant features that may deteriorate the performance of the classification tasks. (2) Omics-specific prediction via GCN. For each omics data type, a weighted patient similarity network was constructed from the omics features. Then, a GCN was trained using both the omics features and the corresponding patient similarity network for omics-specific class prediction. (3) Multi-omics integration via VCDN. A cross-omics discovery tensor was calculated using the initial class predictions from all the omics-specific networks. A VCDN was then trained with the cross-omics discovery tensor to produce final predictions. VCDN can effectively learn the intra-omics and cross-omics label correlations in the higher-level label space for better classification performance with multi-omics data. MORONET is an end-to-end model, where both omics-specific GCN and VCDN are trained jointly. We describe each component in detail in the following sections.

### Pre-processing

To remove noise and experimental artifacts in the data and better interpret the results, proper pre-processing of omics data is essential. First, for DNA methylation data, only probes corresponding to the coding region in Illumina Infinium HumanMethylation27 BeadChip were retained for better interpretability. The number of features for each omics data type is listed in Table 1. Then, we further filtered out features with no signal (zero mean value) and low variances. Specifically, we applied different variance filtering thresholds for different types of omics data: 0.1 for mRNA expression data and 0.001 for DNA methylation data, since different omics data types came with different ranges. For miRNA expression data, we only filtered out features with no variation (variance equals to zero) as the available features in the original datasets were limited due to the small number of miRNAs. The same variance thresholds were used across all classification experiments.

Since each type of omics data could contain redundant features that might have negative effects on the classification performance, we further pre-selected the omics features through statistical tests. For each classification task, ANOVA F-value was calculated sequentially using the training data to evaluate whether a feature was significantly different across different classes. False discovery rate (FDR) - controlling procedures were applied for multiple-testing compensation and the top 200 most significant features for each omics data type were selected. Finally, we individually scaled each type of omics data to [0, 1] through linear transformations for training MORONET.

### GCN for omic-specific learning

We utilized GCN for omic-specific learning in MORONET, where a GCN is learned for each omics data type to perform classification tasks. By viewing each sample as a node in the patient similarity network, the goal of each GCN is to learn a function of features on a graph 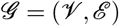 to perform classification tasks by taking advantage of both the features of each node and the relationships between nodes characterized by the graph 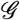. Therefore, a GCN model takes the following two inputs. One input is a feature matrix ***X*** ∈ ℝ^*n*×*d*^, where *n* is the number of nodes and *d* is the number of input features. The other input is a description of the graph structure, which can be represented in the form of an adjacency matrix ***A*** ∈ ℝ^*n*×*n*^. A GCN is built by stacking multiple convolutional layers. Specifically, each layer is defined as:

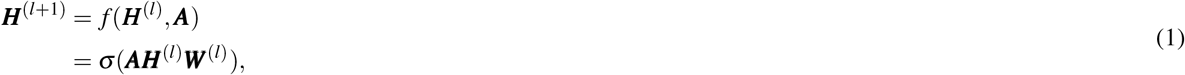

where ***H***^(*l*)^ is the input of the l-th layer and ***W***^(*l*)^ is the weight matrix of the l-th layer. *σ*(·) denotes a non-linear activation function. For effective training of GCN, following the procedure introduced in^58^, we modify the adjacency matrix ***A*** as:

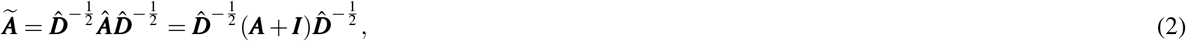

where 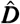 is the diagonal node degree matrix of 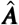 and ***I*** is the identity matrix.

In MORONET, the original adjacency matrix ***A*** is constructed by calculating the cosine similarity between pairs of nodes, and edges with cosine similarity larger than a threshold *ɛ* are retained. Specifically, the adjacency between node *i* and node *j* in the graph, ***A***_*ij*_, is calculated as:

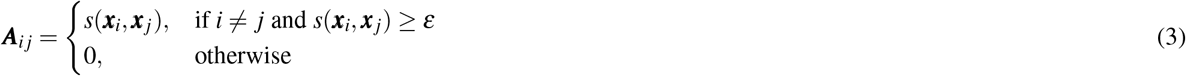

where ***x***_*i*_ and ***x***_*j*_ are the feature vectors of node *i* and node *j*, respectively. 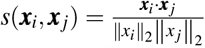 is the cosine similarity between node *i* and *j*. The threshold *ɛ* is determined given a parameter *k*, which represents the average number of edges per node that are retained except self loops:

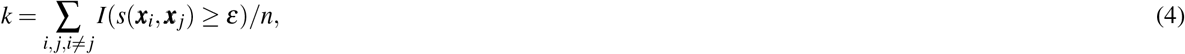

where *I*(·) is the indicator function and *n* is the number of nodes. The parameter *k* for generating the adjacency matrix in Eq. 4 is tuned over {1, 2, 5, 10} through cross validation, and the same *k* value is adopted across all experiments on the same dataset. Although GCN has been widely-utilized in unsupervised^59–62^ and semi-supervised^58, 63–65^ learning, in this paper, we further extend the use of GCN for supervised classification tasks. For training data ***X***_*tr*_ ∈ ℝ^*n*_*tr*_×*d*^, the corresponding adjacency matrix ***A***_*tr*_ ∈ ℝ^*n*_*tr*_×*n*_*tr*_^ can be calculated from Eq. 2. A graph convolutional network *GCN*(·) can be trained on ***X***_*tr*_ and ***A***_*tr*_, where 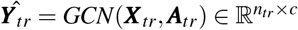 and the *i*-th row of 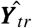 represents the predicted label probability distribution of the *i*-th training sample. *c* denotes the number of classes in the classification task. Therefore, both the features and the graphical structure of the training data are utilized in learning the classification task.

For a new test sample ***x***_*te*_ ∈ ℝ^*d*^, we extend the data matrix to 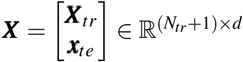 and generate the adjacency matrix to ***A*** ∈ ℝ^(*N*_*tr*_+1)×(*N*_*tr*_+1)^ according to Eq. 2–3. The entries in the last row and last column of ***A*** are the only entries calculated during testing and reflect the affinity between the test sample ***x***_*te*_ and the training samples ***X***_*tr*_. Given ***X***, ***A*** and the trained GCN model *GCN*(·), we have 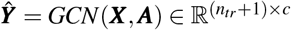. The predicted label probability distribution for the test sample is 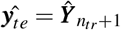, which is the last row of 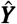. To this end, both the features of the test sample and the correlations between the test sample and the training samples are utilized in predicting the label of the new test sample ***x***_*te*_.

To perform omic-specific classification, for the *i*-th omics data type, we construct a multi-layer GCN denote as *GCN*_*i*_(·) with the output dimensionality of *c*. Given the training data of 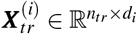 and the corresponding adjacency matrix 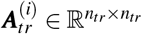 of the *i*-th omics data type, we use L2-norm loss to train the omic-specific GCN:

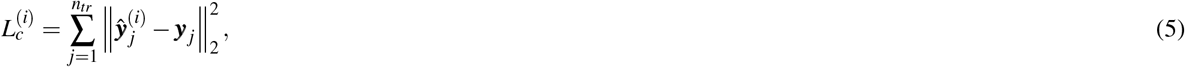

where ***y*** ∈ ℝ^*c*^ is the one-hot encoded label of the *j*-th training sample. 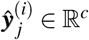 is the predicted label distribution of the *j*-th training sample by *GCN_i_*(·), which is the *j*-th row of the matrix 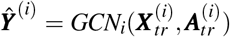. In order to account for the label imbalance in the training data, we further apply different weights on the losses of different classes in Eq. 5, where the weight of a class is set to the inverse of its frequency in the training data.

### VCDN for multi-omics integration

Existing methods utilizing multi-view data on biomedical classification tasks either directly concatenate features from different views, or learn to fuse data from different views either by learning the weights of each view or fusing features from different views in a low-level feature space^4, 66–68^. However, it is always challenging to align various views properly without causing negative influence. On the other hand, VCDN^11^ is designed to learn the higher-level intra-view and cross-view correlations in the label space, and has shown significantly improvements in human action recognition tasks. In MORONET, we utilize VCDN to integrate different omics data types for classification. Moreover, while the original work of VCDN is designed for data with two views^11^, we further extend it to accommodate multiple types of omics data.

Since mRNA expression data, DNA methylation data, and miRNA expression data are utilized in our experiments, for simplicity, we demonstrate how to extend VCDN with three views. Extension to higher number of views can be performed in a similar fashion. For the predicted label distribution of the *j*-th training sample from three different omics data types 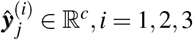, we construct a cross-omics discovery tensor ***C***_*j*_ ∈ ℝ^*c*×*c*×*c*^, where each entry of ***C***_*j*_ is calculated as:

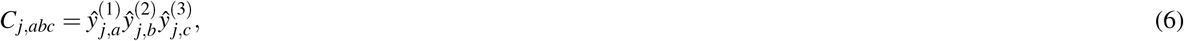

where 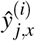 denotes the *x*-th entry of 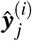. Then, the obtained tensor ***C***_*j*_ is reshaped to a *c*^3^ dimensional vector and forward to *VCDN*(·) for the final prediction. *VCDN*(·) is designed as a two-layer fully-connected network with output dimension of *c* and cross-entropy loss is utilized to train *VCDN*(·):

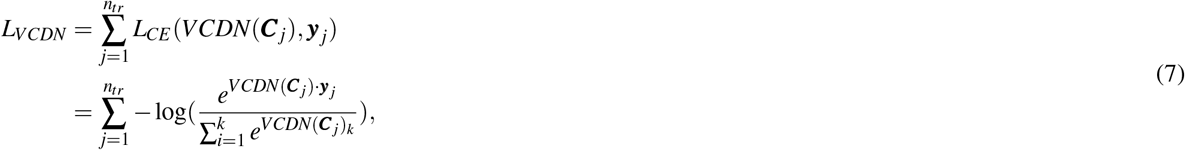

where *L*_*CE*_(·) represents the cross entropy loss function and *VCDN*(***C***_*jk*_) denotes the *k*-th element in the vector *VCDN*(***C***_*j*_) ∈ ℝ^*c*^. To this end, *VCDN*(·) could reveal the latent cross-view label correlations and help to improve the learning performance. By utilizing *VCDN*(·) to integrate initial predictions from different types of omics data, the final prediction made by MORONET is based on both omics-specific predictions and the learned cross-omics label correlation knowledge.

In summary, in our experiments where three omics data types are used, the total loss function of MORONET can be written as:

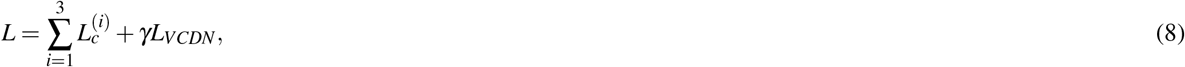

where *γ* is a trade-off parameter between the omics-specific classification loss and the final classification loss from *VCGN*(·). We set *γ* = 1 in all our experiments. MORONET is an end-to-end model and all networks are trained jointly. For training MORONET, during one epoch in the training process, we first fix *VCDN*(·) and update *GCN*_*i*_(·)*, i* = 1, 2, 3 for each omics data type to minimize the loss function *L*. Then we fix the omics-specific GCN and update *VCDN*(·) to minimize *L*. Omics-specific GCN and VCDN are updated alternatively until convergence.

## Availability of data and materials

The ROSMAP dataset was obtained from AMP-AD Knowledge Portal (https://adknowledgeportal.synapse.org/). Omics data of LGG, KIPAN, and BRCA, as well as the grade information of LGG patients were obtained from The Cancer Genome Atlas Program (TCGA) through Broad GDAC Firehose (https://gdac.broadinstitute.org/). PAM50 breast cancer subtypes of TCGA BRCA patients were obtained through TCGAbiolinks R package (http://bioconductor.org/packages/release/bioc/html/TCGAbiolinks.html). The source code of this work can be downloaded from GitHub (https://github.com/txWang/MORONET).

## Acknowledgements

This work was supported by Indiana University Precision Health Initiative and National Institute of Biomedical Imaging and Bioengineering (R01EB025018).

## Author contributions statement

T.W., Z.D., and K.H. conceived and designed the study. T.W. and W.S. performed the computational analysis with assistance from Z.H. T.W., Z.D., and K.H. wrote the manuscript. W.S., Z.H., H.T., and J.Z. edited the manuscript. All the authors reviewed and approved the final manuscript.

